# Altricial bird early-stage embryos express the molecular ‘machinery’ to respond to maternal thyroid hormone cues

**DOI:** 10.1101/2021.12.07.471587

**Authors:** Suvi Ruuskanen, Mikaela Hukkanen, Natacha Garcin, Nina Cossin-Sevrin, Bin-Yan Hsu, Antoine Stier

## Abstract

Maternal hormones, such as thyroid hormones transferred to embryos and eggs, are key signalling pathways to mediate maternal effects. To be able to respond to maternal cues, embryos must express key molecular ‘machinery’ of the hormone pathways, such as enzymes and receptors. While altricial birds begin thyroid hormone (TH) production only at/after hatching, experimental evidence suggests that their phenotype can be influenced by maternal THs deposited in the egg. However, it is not understood, how and when altricial birds express genes in the TH-pathway. For the first time, we measured the expression of key TH-pathway genes in altricial embryos, using two common altricial ecological model species (pied flycatcher, *Ficedula hypoleuca* and blue tit *Cyanistes caeruleus*). Deiodinase *DIO1* gene expression could not be reliably confirmed in either species, but deiodinase enzyme *DIO2* and *DIO3* genes were expressed in both species. Given that DIO2 coverts T4 to biologically active T3, and DIO3 mostly T3 to inactive forms of thyroid hormones, our results suggest that embryos may modulate maternal signals. Thyroid hormone receptor (*THRA and THRB*) and monocarboxyl membrane transporter gene (*SLC15A2*) were also expressed, enabling TH-responses. Our results suggest that early altricial embryos may be able to respond and potentially modulate maternal signals conveyed by thyroid hormones.

## Introduction

Maternal effects are a powerful force shaping offspring phenotype and survival, and may adapt offspring phenotype to a predicted environment (although the adaptiveness is still under debate : Marshall and Uller 2007, Uller et al. 2013, Yin et al. 2019, Sanchez-Tojar et al. 2020, Zhang et al. 2020). Maternal effects can also take different forms, and sometimes bring benefits only to maternal fitness but not to offspring, leading to mother-offspring conflict (Kuijper and Johnston 2018, Groothuis et al. 2020). It has become clear that mechanisms underlying maternal effects are diverse, consisting of nutritional, temperature-related, hormonal, epigenetic, microbe-related and even acoustic signals to the offspring (e.g. Marshall and Uller 2007, Mousseau et al. 2009, DuRant et al. 2013, Groothuis et al. 2019, Mariette et al. 2021). Yet, it is increasingly acknowledged that offspring may not just be passive recipients of the signal, but may actively modify the signal, for example metabolizing maternal hormones, such as steroids (e.g. Paitz et al. 2011, Vassallo et al. 2014, Groothuis et al. 2019, Kumar et al. 2019, Paitz et al. 2020), influencing the resolution of potential parent-offspring conflict.

Thyroid hormones, thyroxine (T4) and biologically active triiodothyronine (T3), are key maternal hormones which critically influence early development in many organisms (e.g. McNabb and Darras 2015). For example, the influence of maternal thyroid hormones on amphibian development was established already in the 1910’s (Gudernatsch 1912), and their importance on human neurodevelopment has been heavily investigated (e.g. Patel et al. 2011). However, the role of maternal (prenatal) thyroid hormones in other systems, such as birds, has not been thoroughly studied until very recently (Ruuskanen and Hsu 2018, Darras 2019, Sarraude et al. 2020b, Sarraude et al. 2020c, d, Stier et al. 2020).

Thyroid hormones of maternal origin are found in eggs of both precocial birds (species with advanced embryonic development and independence after hatching) and altricial birds (species not independent after hatching, Ruuskanen and Hsu 2018). To be able to respond to maternal thyroid hormone signalling, embryos must have the molecular ‘machinery ’ of the thyroid axis (TH-axis) in place: they need to express for example transmembrane transporters (e.g. monocarboxyl membrane transporters) transporting hormones to cells, cellular deiodinases, which convert T4 to bioactive T3 and to inactive forms (rT3 and T2), and intracellular hormone receptors (THRA and THRB, Zoeller et al. 2007). Embryos of precocial birds have been discovered to contain thyroid hormones and express genes in the TH-axis, such as *DIO2* as early as 4 days into incubation (Van Herck et al. 2012). The expression also varied depending on maternal hormonal concentrations (Van Herck et al. 2012). Importantly, precocial birds begin embryonic thyroid production around mid-incubation while in contrast, altricial birds are only able to produce thyroid hormones at/after hatching (Darras 2019), thus being potentially dependent on maternal hormones during the entire embryonic period. Thyroid hormones (likely of maternal origin) were indeed shown to be present in embryonic plasma of altricial species such as ring dove (*Streptopelia risoria*) and European starling (*Sturnus vulgaris*) before the presumed timing of thyroid gland maturation (McNabb and Cheng 1985, Schew et al. 1996). Furthermore, it has been recently experimentally shown that egg thyroid hormones in altricial species can influence pre-and post-hatching development, such as embryo survival, growth and physiology (Ruuskanen et al. 2016, Hsu et al. 2019, 2020, Sarraude et al. 2020a, Sarraude et al. 2020b, Stier et al. 2020). It is not however understood if, how and when altricial species express genes of thyroid hormone response ‘machinery’, whereby maternal hormonal effects could take place.

The aim of the study was to characterize expression of thyroid hormone signalling-related genes in early development of altricial birds. To this end, we collected early embryos of different ages from two common altricial species often used as models in ecological and evolutionary research, the pied flycatcher (*Ficedula hypoleuca*), and the blue tit (*Cyanistes caeruleus*). We measured expression of key thyroid-related genes (1) a membrane transporter (*SLC15A2*), (2) deiodinases (*DIO1-3*), and (3) thyroid hormone receptors (*THRA* and *THRB*). We characterized the expression of the selected genes across embryos of different age to reveal potential age-related changes. The characterization of the gene expression allows us to understand when and how altricial birds may respond to maternal thyroid hormone cues. Furthermore, expression of DIOs would also indicate that early embryos may be capable of metabolizing maternal hormones, potentially modulating maternal signalling.

## Material and methods

Sample collection was in accordance with all relevant guidelines and regulations and approved by the Environmental Center of Southwestern Finland (license number VARELY924/2019). The data collection was conducted in spring-summer 2020 in nest box population of blue tits and pied flycatchers in south-western Finland (60° 25⍰ N, 22° 10⍰ E). We monitored the population for the initiation of egg laying, marked each egg in laying order, and visited the nest daily to record the start of incubation. We collected one egg from 10 nests per species to limit consequences on their breeding success. The collected egg was positioned at middle of the laying order to avoid any laying-order associated variation as reported for egg composition, especially for first and last eggs (e.g. Hsu et al. 2019). The collected eggs were kept warm until dissection (within 1-2h). The embryo was carefully removed from the yolk (using equipment treated with RNase decontamination solution, RNase*Zap*®, ThermoFischer), immediately frozen in liquid nitrogen and stored at -80ºC for ca. 5 months. The eggs varied in the duration of incubation and embryos were staged based on Murray et al. (2013) with 0.5 day accuracy.

We analysed expression levels of six genes of interest using RT-qPCR. These included a monocarboxyl membrane transporter (*SLC15A2*), all three deiodinases (*DIO1, DIO2, DIO3*) and thyroid hormone receptor genes (*THRA, THRB*). Reference genes were selected from prior publications in blue tits (Capilla-Lasheras et al. 2017). Primers for reference genes were designed on exon-exon junction using NCBI primer blast (Table 1). Initially four reference genes (*ACTB, GADPH, SDHA* and *TRFC*) were selected for validation.

**Table 1.**
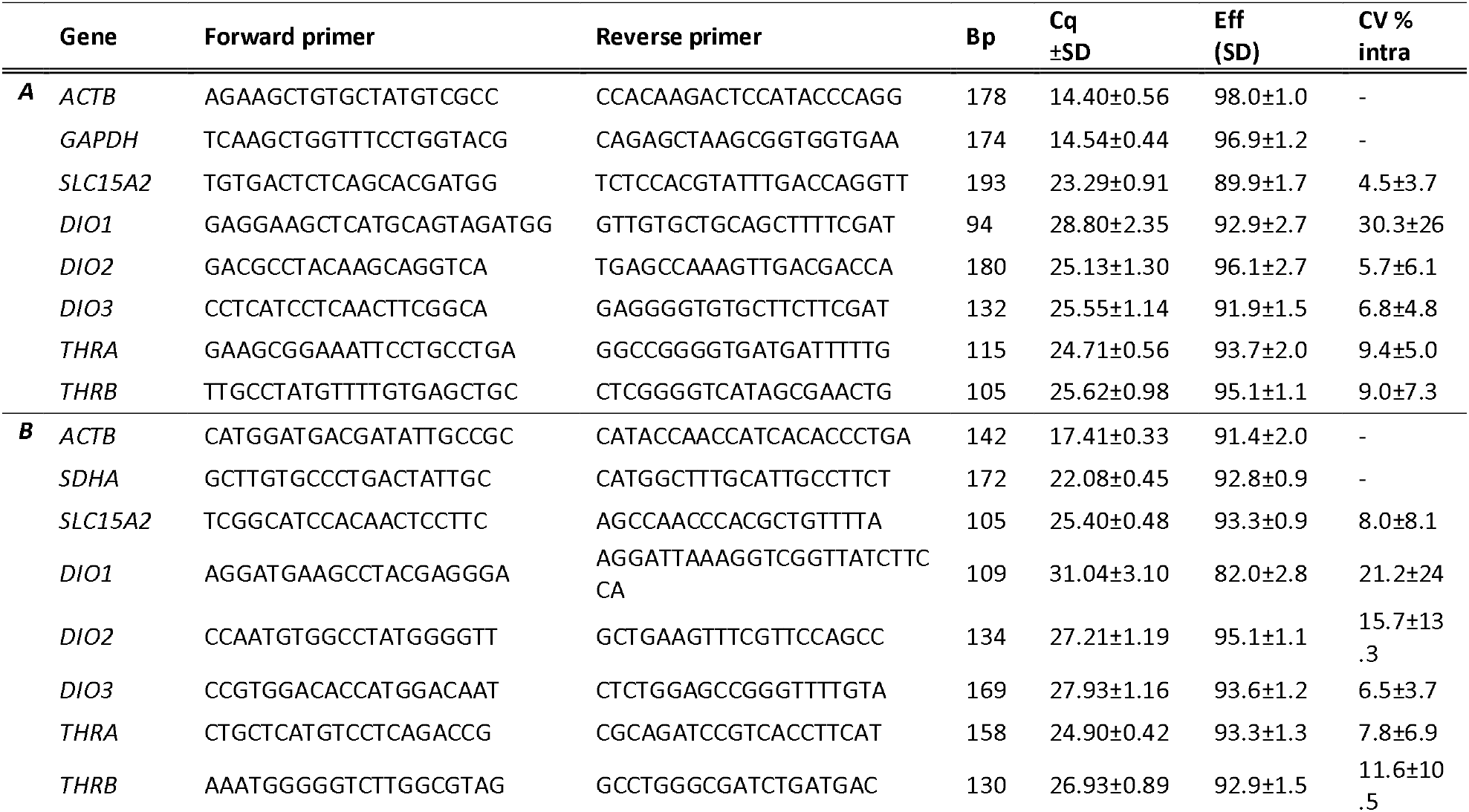
Forward and reverse primer sequences for refence and target genes for (A) blue tit and (B) pied flycatcher (from collared flycatcher genome). Cq refers to qPCR quantitation cycle (a higher value indicating a lower initial target mRNA amount), efficiency has been evaluated using LinReg method and technical precision estimated as coefficient of variation (CV in %) for final ratios at the intra-plate level (based on duplicates). All the samples for one gene per species were run on one single plate.

RNA was extracted from whole-embryos using Nucleospin RNA Plus extraction kit (Macherey-Nagel), following manufacturer’s instructions and stored at -80ºC for 2 months. RNA concentration and purity were quantified using optical density. Samples not meeting quality criteria ([RNA] > 30 ng/μl, 260/280 and 260/230 > 1.80) were excluded for further analysis. RNA integrity was checked using E-Gel 2% electrophoresis system (Invitrogen), and the ribosomal RNA 18S vs. 28S bands intensity, and deemed satisfactory. 500ng of RNA were used for cDNA synthesis using the SensiFAST cDNA Synthesis kit (Bioline) following manufacturer instructions. cDNA was diluted at a final concentration of 1.2 ng/μl for qPCR analysis. No-RT control samples were prepared following the same protocol, but without reverse transcriptase enzyme.

Primers for the target genes (see Table 1) were designed using NCBI primer blast, to exon-exon junction whenever possible. Blue tit reference genome was assembly <u>GCA_002901205.1</u>. For pied flycatcher, the reference genome was not available and thus the genome of a closely related species, the collared flycatcher genome was used (assembly <u>GCA_000247815.2</u>). To validate the primers, initially 2-5 primers for each gene were designed and tested for specificity, efficiency and linearity. Pooled samples (pooling RNA from three individuals) from both species were used in validation. Specificity was checked using BLAST analysis and confirmed by a single narrow peak in melting curve analyses and the presence of a single PCR product of the expected size on agarose gel. Amplification of controls with no reverse transcriptase never occurred before at least 7 cycles after the lower Cq sample (except for *DIO1* that was excluded from interpretation, see below), and thus contamination by genomic DNA could not interfere with our results. Based on their performance during initial validation, *ACTB* (actin beta, highly conserved protein involved e.g in cell motility) and *GAPHD* (glyceraldehyde-3-phosphate dehydrogenase, a key protein in carbo-hydrate metabolism) were used for reference genes for blue tit gene expression, and *ACTB* and *SDHA* (succinate dehydrogenase complex flavoprotein subunit A, a key mitochondrial protein) for pied flycatchers.

Samples and controls (two controls per plate) were analysed in duplicates. All samples for one gene were run in one plate, and two genes were analysed per qPCR plate. qPCR was performed in a total volume of 12μl containing 5μl of each diluted cDNA sample (i.e. 1.2ng/μl) and 7μl of reaction mix containing primers (forward and reverse) at a final concentration of 300nM and Sensifast SYBR®Low-ROX Mix (Bioline). qPCR assays were performed on a Mic qPCR instrument (Bio Molecular Systems) and included a two-step cycling with the following conditions: 2 minutes at 95°C; then 40 cycles of 5s at 95°C followed by 20s at 60°C (fluorescence reading) for all reactions. The expression of each gene was calculated as (1+Ef_Target_)^ΔCq(Target)^ / geometric mean [(1+Ef_Ref_gene1_)^ΔCq(Ref_gene1)^ + (1+Ef _Ref_gene2_)^ΔCq(Ref_gene2)^], Ef being the amplification’s efficiency and ΔCq being the difference between the Cq-values of the reference sample and the sample of interest. Statistical analyses were not conducted because of the limited number of replicates.

## Results

In both species, pied flycatchers and blue tits, the coefficients of variation for DIO1 were high (many samples with >30%) and Cq values also high (being < 5 cycles apart from no-RT controls) and therefore its expression could not be reliably measured. All other genes, membrane transporters (*SCL152A*), deiodinase enzymes (*DIO2, DIO3*) and TH receptors (*THRA and THRB* subunits) were expressed in both altricial species, but at relatively low levels compared to reference genes (Cq of target genes being << reference genes; Table 1). None of the genes showed clear changes with embryonic development time (Fig 1). Yet, for DIO2 the expression levels of older (4-5-day old embryos) seemed to be higher, and there were specifically some individuals with high expression values especially the oldest (5.5 days) embryos. When visually inspecting the expression patters for both species, few embryos sampled from the earliest time-points (1-day old embryos) showed somewhat high expression for part of the genes (*DIO3, THRA, THRB*) compared to other time-points.

**Fig 1.**
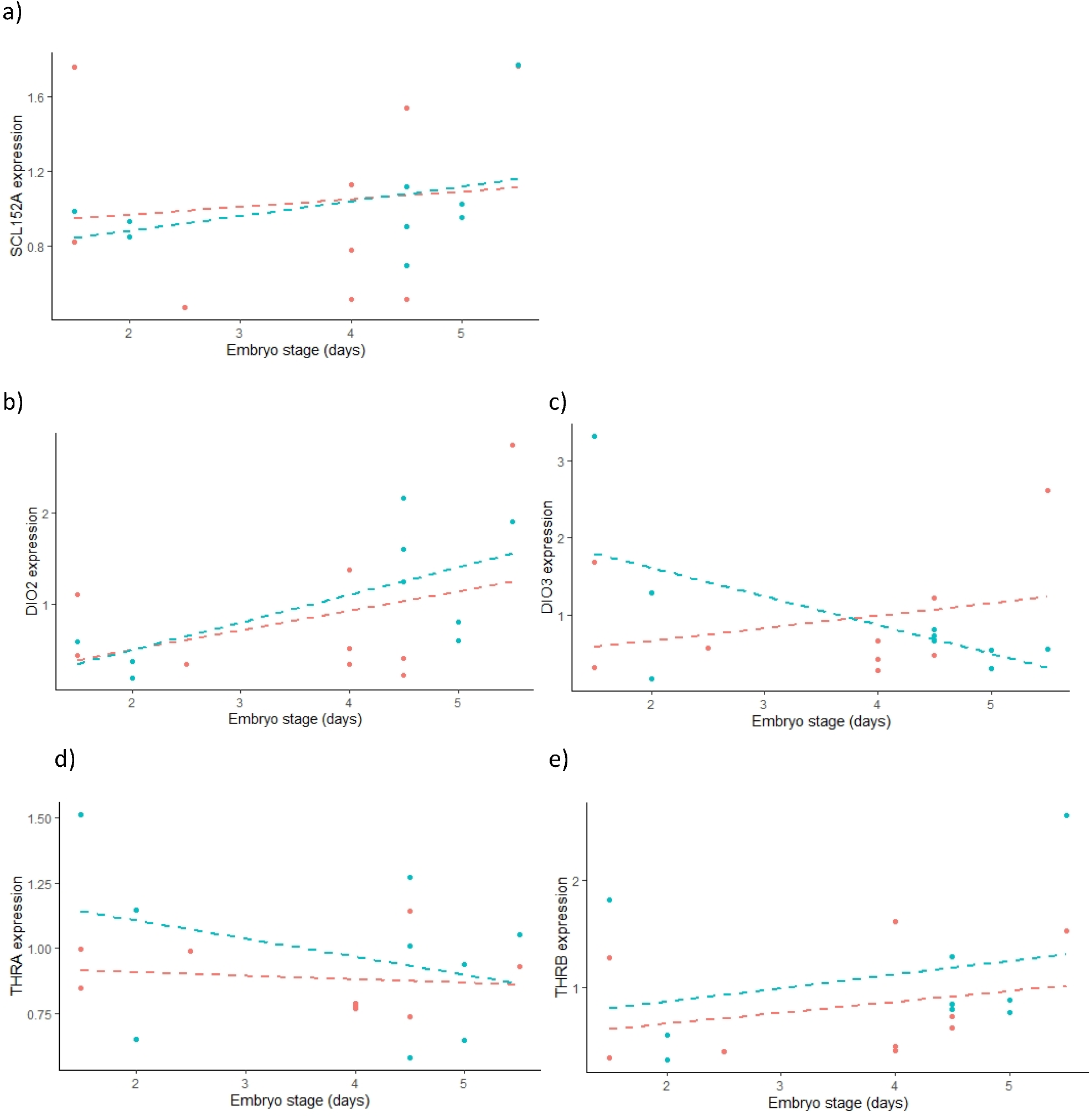
Expression of thyroid hormone axis related genes in embryos (1-5.5 days) of the altricial pied flycatcher (blue) and blue tit (red): (A) membrane transporter (*SLC15A2*), (B) deiodinase 2 (DIO2) converting T4 to T3, (C) deiodinase 3 (DIO3) converting T4 and T3 to the inactive form rT3, (D) thyroid hormone receptor A (THRA) & (E) thyroid hormone receptor B (THRB). N = 10 individuals per species. Dashed lines are included for visualization, but due to low sample sizes, statistical analyses have not been performed. Species cannot be directly compared as relative gene expression was evaluated in a species-specific manner (i.e. different primers and reference genes).

## Discussion

We were able to detect expression of the deiodinase enzyme genes *DIO2* and *DIO3* in early altricial embryos of two passerine species, blue tits and pied flycatchers. *DIO1* could not be reliably measured in either species. DIO1 is mostly a scavenger enzyme, converting inactive rT3 to other inactive forms (e.g. Darras et al. 2019). In previous studies in 4-day-old precocial chicken (*Gallus gallus domesticus*), *DIO1* was expressed but did not yield a functional enzyme (Van Herck et al. 2012). DIO2, in turn, is the key enzyme converting T4 to the active form T3. Therefore, expression of this enzyme early in prenatal development, along with our findings of expression of transmembrane transporter gene (*SCL152A*) and thyroid hormone receptor genes (*THRA and THRB*) in altricial embryos would support the hypothesis that altricial embryos may respond to maternal thyroid hormones before their own thyroid hormone production. In precocial birds, *DIO2* gene expression increased during embryonic development (from day 4 onwards), whereas *DIO3* gene expression was more variable and cell-type dependent (Geysens et al. 2012, Van Herck et al. 2012). Interestingly, DIO3 mainly converts T3 to inactive forms, and its expression can be seen as regulating the cellular exposure to active T3. Given that mothers deposit also T3 into egg yolks, expression of DIO3 in the early embryo would open up the possibility that embryos can downregulate maternal signalling, as observed for androgen hormones (reviewed in Groothuis et al. 2019). A further validation step would include verifying the translation of these transcripts to functional proteins, e.g. using western blots or proteomic approaches.

In our data, few samples from earliest time-points (ca 1-day-old embryo), seemed to show rather high expression levels for some genes. In other taxa, such as fish embryos, transcripts in very early embryos are predicted to be of maternal origin (e.g. Essner et al. 1997, Takayma et al. 2008). For example, Vergauwen et al. (2018) confirmed the presence of maternal transfer of TPO (thyroid peroxidase), DIO1-3, THRA and THRB mRNA using unfertilized eggs, yet levels quickly decreased during embryo development. Maternal mRNA transfer has rarely been explored in birds beyond studies related to fertilization (Olszanska and Stepinska 2008) and to our knowledge there is no data on maternal thyroid hormone related mRNAs in eggs. Thus, it would be important to verify if and how much of the transcripts may be of maternal origin, by sampling unincubated (and preferably unfertilized) eggs across species. Yet, there are technical challenges in working with low levels of transcripts in lipid-rich yolk tissue, especially for (wild) species with small eggs.

All in all, thyroid hormone signalling and its consequences on early development in (altricial) birds is a fruitful avenue for further research. Knowledge gained from early-life thyroid-related gene expression is not only important from the perspective of fundamental developmental biology and comparative physiology, but also for (eco)toxicology: wild bird species are subject to various endocrine disrupting chemicals (EDCs) also via the egg (e.g. Ruuskanen et al. 2014). Thyroid disruption via EDCs can occur at multiple locations within the thyroid axis, acting through several molecular targets, such as inhibition of T4 production, inhibition of deiodination of T4 to T3 in peripheral tissues, and impacts on TH receptors (McNabb 2007). Identifying molecular targets (when and how embryos respond to THs) could help in understand and screening for the prenatal effects of EDCs.

## Acknowledgements

We thank volunteer helpers for monitoring the population. This study was financially supported by the Academy of Finland (#286278 to SR) and the Turku Collegium for Science and Medicine (grants to SR and AS). NCS acknowledges support from the EDUFI Fellowship and Maupertuis Grant. B-Y.H work was supported by a grant from the Ella and Georg Ehrnrooth Foundation. AS was supported by a Marie Sklodowska-Curie Postdoctoral Fellowship (#894963) at the time of writing.

## Author contributions

SR, AS and BYH conceived the study. SR, MH, AS and NC contributed to data collection. SR, AS and NG conducted the laboratory analyses. SR prepared the first draft and all authors commented on the draft.

## Notes

### Competing Interest Statement

The authors have declared no competing interest.

## References

Capilla-Lasheras, P., D. M. Dominoni, S. A. Babayan, P. J. O’Shaughnessy, M. Mladenova, L. Woodford, C. J. Pollock, T. Barr, F. Baldini, and B. Helm. 2017. Elevated Immune Gene Expression Is Associated with Poor Reproductive Success of Urban Blue Tits. Frontiers in Ecology and Evolution 5:13.

Darras, V. M. 2019. The Role of Maternal Thyroid Hormones in Avian Embryonic Development. Frontiers in Endocrinology 10:10.

DuRant, S. E., Hopkins, W. A., Hepp, G. R., & Walters, J. R. 2013. Ecological, evolutionary, and conservation implications of incubation temperature□dependent phenotypes in birds. Biological Reviews, 88(2), 499–509.

Essner J.J., et al., 1997 The zebrafish thyroid hormone receptor α1 is expressed during early embryogenesis and can function in transcriptional repression. Differentiation 62, 107–117.

Geysens, S., J. L. Ferran, S. L. J. Van Herck, P. Tylzanowski, L. Puelles, and V. M. Darras. 2012. DYNAMIC mRNA DISTRIBUTION PATTERN OF THYROID HORMONE TRANSPORTERS AND DEIODINASES DURING EARLY EMBRYONIC CHICKEN BRAIN DEVELOPMENT. Neuroscience 221:69–85.

Groothuis, T. G. G., B. Y. Hsu, N. Kumar, and B. Tschirren. 2019. Revisiting mechanisms and functions of prenatal hormone-mediated maternal effects using avian species as a model. Philosophical Transactions of the Royal Society B-Biological Sciences 374:9.

Groothuis, T.G.G., Kumar, N., Hsu, B-Y. 2020. Explaining discrepancies in the study of maternal effects: the role of context and embryo. Current Opinion Behavioral Science 36: 185–192.

Gudernatsch, J.F. 2012. Feeding Experiments on tadpoles. I. The intlnence of specific organs given as food on growth and differentiation. A contribution to the knowledge of organs with internal secretion. ARCHIV FUR ENTWICKLUNGSMECHANIK DER ORGANISMEN. 35, 457–483.

Hsu, B-Y., Verhagen, I., Gienapp, P., Darras, V., Visser, M. 2019. Between-and Within-Individual Variation of Maternal Thyroid Hormone Deposition in Wild Great Tits (Parus major). American Naturalist 194, E96–108.

Hsu, B. Y., T. Sarraude, N. Cossin-Sevrin, M. Crombecque, A. Stier, and S. Ruuskanen. 2020. Testing for context-dependent effects of prenatal thyroid hormones on offspring survival and physiology: an experimental temperature manipulation. Scientific Reports 10.

Hsu, B.-Y., Doligez, B., Gustafsson, L. & Ruuskanen, S. 2019 Transient growth-enhancing effects of elevated maternal thyroid hormones at no apparent oxidative cost during early postnatal period. J. Avian Biol. 50, jav–01919.

Kuijper, B., Johnston, R. 2018. Maternal effects and parent–offspring conflict. Evolution 72: 220–233.

Kumar, N., A. van Dam, H. Permentier, M. van Faassen, I. Kema, M. Gahr, and T. G. G. Groothuis. 2019. Avian yolk androgens are metabolized rather than taken up by the embryo during the first days of incubation. Journal of Experimental Biology 222:6.

Mariette, M. M., Clayton, D. F., & Buchanan, K. L. 2021. Acoustic developmental programming: a mechanistic and evolutionary framework. Trends in Ecology & Evolution.

Marshall, D. J., and T. Uller. 2007. When is a maternal effect adaptive? Oikos 116:1957–1963.

McNabb, F. M. A. 2007. The hypothalamic-pituitary-thyroid (HPT) axis in birds and its role in bird development and reproduction. Critical Reviews in Toxicology 37:163–193.

McNabb, F. M. A., and M. F. Cheng. 1985. THYROID DEVELOPMENT IN ALTRICIAL RING DOVES, STREPTOPELIA-RISORIA. General and Comparative Endocrinology 58:243–251.

Mousseau, T., Uller, T., Wapstra, E., Badyaev, A.V. 2009. Evolution of maternal effects: past and present. Phil. Trans. R. Soc. B 2009 364, 1035–1038.

Murray, J. R., C. W. Varian-Ramos, Z. S. Welch, and M. S. Saha. 2013. Embryological staging of the Zebra Finch, Taeniopygia guttata. Journal of Morphology 274:1090–1110.

Olszanska, B., and U. Stepinska. 2008. Molecular aspects of avian oogenesis and fertilisation. International Journal of Developmental Biology 52:187-194.’

Paitz, R.T., Bowden, R.M., Castro, J.M., 2011. Embryonic modulation of maternal steroids in European starlings (Sturnus vulgaris). Proceedings of the Royal Society B: Biological Sciences 278 (1702), 99–106

Paitz, R.T., Angles, R., Cagnev, E., 2020. In ovo metabolism of estradiol to estrone sulfate in chicken eggs: implications for how yolk estradiol influences embryonic development. General and Comparative Endocrinology 287, 113320.

Patel J., K. Landers, H. Li, R.H. Mortimer, and K. Richard. 2011. Thyroid hormones and fetal neurological development. J Endocrinol 209:1–8.

Ruuskanen, S., V. M. Darras, M. E. Visser, and T. G. G. Groothuis. 2016. Effects of experimentally manipulated yolk thyroid hormone levels on offspring development in a wild bird species. Hormones and Behavior 81:38–44.

Ruuskanen, S., and B. Y. Hsu. 2018. Maternal Thyroid Hormones: An Unexplored Mechanism Underlying Maternal Effects in an Ecological Framework. Physiological and Biochemical Zoology 91:904–916.

Ruuskanen, S., T. Laaksonen, J. Morales, J. Moreno, R. Mateo, E. Belskii, A. Bushuev, A. Jarvinen, A. Kerimov, I. Krams, C. Morosinotto, R. Maend, M. Orell, A. Qvarnstrom, F. Slater, V. Tilgar, M. E. Visser, W. Winkel, H. Zang, and T. Eeva. 2014. Large-scale geographical variation in eggshell metal and calcium content in a passerine bird (Ficedula hypoleuca). Environmental Science and Pollution Research 21:3304–3317.

Sanchez-Tojar, A., M. Lagisz, N. P. Moran, S. Nakagawa, D. W. A. Noble, and K. Reinhold. 2020. The jury is still out regarding the generality of adaptive ‘transgenerational’ effects. Ecology Letters 23:1715–1718.

Sarraude, T., B.-Y. Hsu, T. Groothuis, and S. Ruuskanen. 2020a. Manipulation of prenatal thyroid hormones does not influence growth or physiology in nestling pied flycatchers. Physiological and Biochemical Zoology.

Sarraude, T., B. Y. Hsu, T. G. G. Groothuis, and S. Ruuskanen. 2020b. Manipulation of Prenatal Thyroid Hormones Does Not Affect Growth or Physiology in Nestling Pied Flycatchers. Physiological and Biochemical Zoology 93:255–266.

Sarraude, T., B. Y. Hsu, T. G. G. Groothuis, and S. Ruuskanen. 2020c. Testing different forms of regulation of yolk thyroid hormone transfer in pied flycatchers. Journal of Experimental Biology 223.

Schew, W. A., F. M. A. McNabb, and C. G. Scanes. 1996. Comparison of the ontogenesis of thyroid hormones, growth hormone, and insulin-like growth factor-I in ad Libitum and food-restricted (altricial) European starlings and (precocial) Japanese quail. General and Comparative Endocrinology 101:304–316.

Stier, A., Bize, P., Hsu, B.-Y. & Ruuskanen, S. 2019 Plastic but repeatable: rapid adjustments of 239 mitochondrial function and density during reproduction in a wild bird species. Biol Letters 15, 20190536.

Stier, A., B. Y. Hsu, C. Marciau, B. Doligez, L. Gustafsson, P. Bize, and S. Ruuskanen. 2020. Born to be young? Prenatal thyroid hormones increase early-life telomere length in wild collared flycatchers. Biology Letters 16:4.

Takayama S, et al., 2008. An F-domain introduced by alternative splicing regulates activity of the zebrafish thyroid hormone receptor α: role of zebrafish TRα F-domain. General and Comparative Endocrinology 155, 176–189.

Uller, T., Nakagawa, S., English, S. 2013. Weak evidence for anticipatory parental effects in plants and animals. Journal of Evolutionary Biology 26: 2161–2170

Van Herck, S. L. J., S. Geysens, J. Delbaere, P. Tylzanowski, and V. M. Darras. 2012. Expression profile and thyroid hormone responsiveness of transporters and deiodinases in early embryonic chicken brain development. Molecular and Cellular Endocrinology 349:289–297.

Vassallo, B.G., Paitz, R.T., Fasanello, V.J., Haussmann, M.F., 2014. Glucocorticoid metabolism in the in ovo environment modulates exposure to maternal corticosterone in Japanese quail embryos (Coturnix japonica). Biology letters 10 (11), 20140502

Vergauwen, L., J. E. Cavallin, G. T. Ankley, C. Bars, I. J. Gabriels, E. D. G. Michiels, K. R. Fitzpatrick, J. Periz-Stanacev, E. C. Randolph, S. L. Robinson, T. W. Saari, A. L. Schroeder, E. Stinckens, J. Swintek, S. J. Van Cruchten, E. Verbueken, D. L. Villeneuve, and D. Knapen. 2018. Gene transcription ontogeny of hypothalamic-pituitary-thyroid axis development in early-life stage fathead minnow and zebrafish. General and Comparative Endocrinology 266:87–100.

Yin, J. J., M. Zhou, Z. R. Lin, Q. S. Q. Li, and Y. Y. Zhang. 2019. Transgenerational effects benefit offspring across diverse environments: a meta-analysis in plants and animals. Ecology Letters 22:1976–1986.

Zhang, Y. Y., J. J. Yin, M. Zhou, Z. R. Lin, and Q. S. Q. Li. 2020. Adaptive transgenerational effects remain significant. Ecology Letters 23:1719–1720.

Zoeller, R. T., S. W. Tan, and R. W. Tyl. 2007. General background on the hypothalamic-pituitary-thyroid (HPT) axis. Critical Reviews in Toxicology 37:11–53.

